# Threonine Supplementation Reduces Methylglyoxal Overflow by Increasing Glycolysis Flux at the Payoff Phase: A Metabolic Modeling Analysis

**DOI:** 10.64898/2026.09.17.751990

**Authors:** Sandra Francis, Rahad Rehas, Yuttamol Muangkram, Niyas Rehman, Anoop Kumar G Velikkakath, Shuhei Noda, Michihiro Araki

**Author notes:** Equal contribution (Sandra Francis, Rahad Rehas, Yuttamol Muangkram).

## Abstract

Methylglyoxal (MGO) is a metabolic byproduct of sugar metabolism involved in the formation of advanced glycation end products (AGEs) and its accumulation disrupts protein function, redox balance and cellular viability. Yet the metabolic rewiring required to divert MGO overflow and prevent these cytotoxic effects remains largely unknown. To address these gaps, we used flux sampling and flux balance analysis in genome scale metabolic model iJO1366 of *E. coli*. High glucose increased MGO flux from 0.16 mmol/gDCW/h to 1.87 mmol/gDCW/h in simulation. Large scale *in silico* screening of metabolic reactions identified that increasing threonine uptake, thereby increasing ethanol flux, reduced glucose induced increase in MGO flux from 1.87 mmol/gDCW/h to 0.405 mmol/gDCW/h. *In silico* inhibition of ethanol production (NAD^+^ regeneration) inhibited the MGO lowering potential of threonine. Threonine supplementation actively drives the acetaldehyde to ethanol flux to provide localized NAD^+^ relief, which subsequently enhances the flux of payoff phase in glycolysis (glyceraldehyde 3-phosphate), thereby efficiently draining the stagnated DHAP pool and shutting down the overflow towards MGO production. Unexpectedly, under reduced oxygen conditions, high glucose caused only a minimal increase in MGO flux, which was not reduced by threonine supplementation. Our study also decoded the mechanistic details behind this paradox. Though this model driven hypothesis requires further validation *in vivo,* these stoichiometric predictions have applications in metabolic engineering, for commercial MGO production. Furthermore, *in vivo* studies focused on bacterial stress, gut microbiome imbalances and AGE related diseases might benefit from these simulations.

**Figure Abstract:** 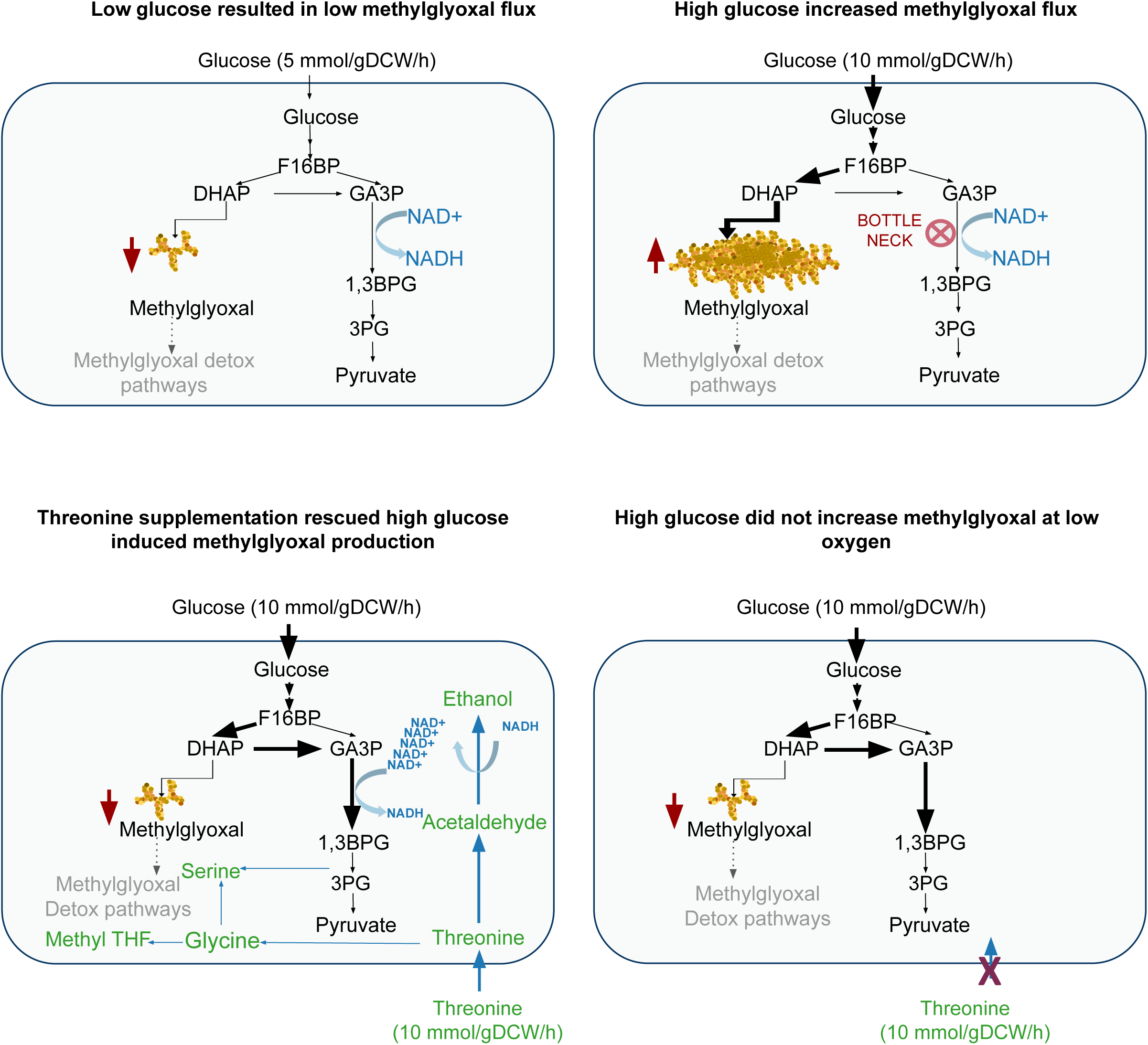

## 1. Introduction

Methylglyoxal (MGO) is a highly reactive α-ketoaldehyde generated as a glycolytic overflow metabolite and serves as a central precursor in the formation of advanced glycation end products (AGEs) (1). As a potent electrophile, MGO readily modifies proteins, nucleic acids and lipids (2) making its intracellular abundance a critical determinant of cellular homeostasis. The reported cytotoxicity of MGO is cleared using several detoxification mechanisms like the glyoxalase system. The glyoxalase system utilizes glutathione and the enzymes (Glyoxalase) GLO1 and GLO2 to convert over 99% of MGO into harmless D-lactate. When this pathway is overwhelmed or glutathione is depleted, backup systems like DJ-1 (PARK7), AKRs (Aldo-Keto Reductases) and ALDHs (Aldehyde Dehydrogenases) step in to neutralize the remaining toxin and protect the cell from damage (3, 4, 5).This entire process is energy consuming and hence can shift the metabolic balance of the system (6).

Maillard autoxidation of hexoses and unsaturated fatty acids, dehydration of dihydroxyacetone could also lead to the formation of MGO as an endogenous route of entry to the metabolism. Although MGO is produced across a wide spectrum of physiological and stress related conditions (7), healthy cells maintain MGO at exceptionally low basal concentrations through stringent metabolic regulation (8). Even modest perturbations to this balance can be catastrophic, millimolar MGO levels arrest growth and trigger widespread cytotoxicity (9).

In *E. coli*, MGO accumulation has long been observed during aerobic growth on glycerol (10, 11), underscoring the sensitivity of central carbon metabolism to redox and energy imbalances. The dominant bacterial source of MGO is the conversion of dihydroxyacetone phosphate (DHAP) by methylglyoxal synthase (EC 4.2.99.11) (12), a reaction that bypasses the tightly regulated lower glycolytic steps. This “pressure-release” shunt provides a rapid means of diverting carbon when glycolytic flux becomes saturated or when NADH/NAD^+^ ratios are perturbed, but it also exposes the cell to the risk of MGO overload.

In humans, genetic analyses demonstrate that triosephosphate isomerase deficiency increases MGO production through impaired interconversion of DHAP and glyceraldehyde 3-phosphate (13). Whereas in *E. coli* deletion of *mgsA* (strain LY168) redirects central carbon flux, suppresses methylglyoxal formation, and enables simultaneous glucose-xylose co-utilization, resulting in increased ethanol production from mixed sugars (14). These phenotypes suggest that MGO metabolism has broader implications for carbon partitioning, redox balance and overflow pathways than previously appreciated across a diverse range of organisms.

Yet despite decades of biochemical, genetic and physiological study, the endogenous mechanisms that buffer, redirect, or dissipate MGO overflow remain incompletely defined. While the glyoxalase system and several reductive detoxification routes are known, the upstream metabolic strategies that prevent MGO accumulation, particularly those capable of physiologically reducing MGO flux remain poorly characterized. This gap limits our understanding of how cells dynamically manage glycolytic stress and constrains efforts to engineer microbial systems with improved robustness or optimized carbon utilization.

The objective of our study is to identify metabolic strategies that can modulate MGO overflow in *E. coli*. Preventing MGO formation may reduce the cellular burden and help preserve metabolic and energy resources that would otherwise be directed towards detoxification of these toxic metabolites. To study this, we used genome scale metabolic models, flux balance analysis (FBA) and flux sampling to systematically examine metabolic states associated with MGO overflow and identify perturbations capable of reducing MGO production. The resulting interventions may provide insights into how *E. coli* maintains metabolic balance under conditions of carbon and redox stress. Thus, this work offers potential approaches for engineering *E. coli* with strategies to reduce accumulation of toxic secondary metabolites during industrial fermentation of products and even for commercial production of MGO itself, which is considered a strong antibiotic (15).

## 2. Materials and Methods

### 2.1 Metabolic Model and Software Environment

All constraint-based modelling simulations were performed in Python 3 using the COBRApy modeling framework (16). The *E. coli* K-12 MG1655 reconstruction iJO1366 (17) served as the reference model and was used in SBML format without modification. The iJO1366 model is composed of 1,366 genes, 2,583 reactions, and 1,805 metabolites. Linear optimization was executed using COBRApy’s default GLPK solver. Data processing, statistical analysis, and visualization were performed using pandas, SciPy, matplotlib and seaborn. The workflow of the simulation and analysis, using the metabolic model as a starting point, is shown in (Figure 1).

**Figure 1.**
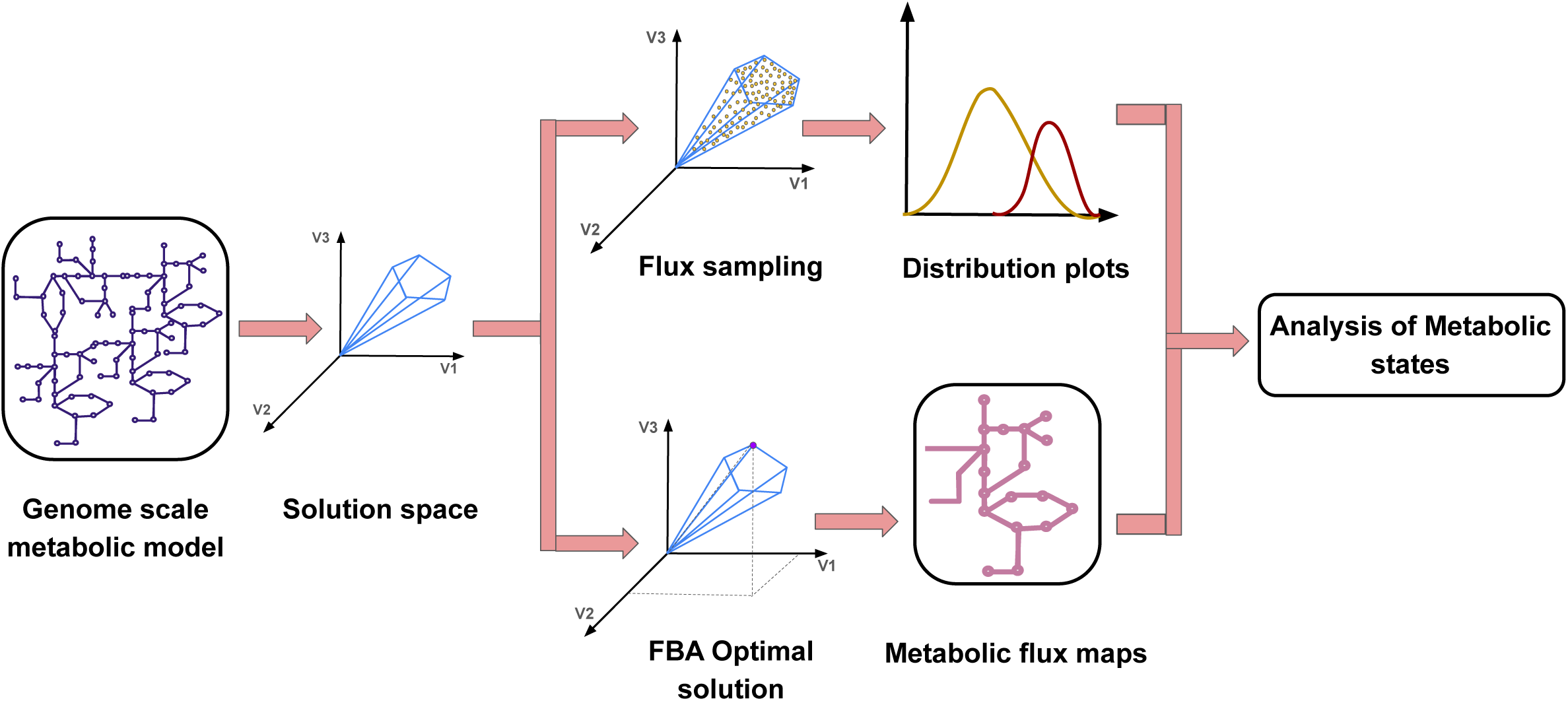
Workflow diagram from the genome scale metabolic model into analysis of metabolic states using flux sampling and FBA.

### 2.2 Flux Sampling

Flux sampling was used to characterize the feasible steady state flux space of the metabolic network by generating probability distributions of reaction fluxes across high dimensional solution states (18). This approach enables systematic exploration of alternative metabolic configurations under defined environmental and genetic constraints (19). Flux sampling was performed in the *E. coli* genome scale metabolic model iJO1366 (17).

#### 2.2.1 Amino Acid Supplementation Screening

We performed a systematic screen of all 20 amino acids to identify potential modulators of MGSA flux. Four fixed nutrient environments were examined, glucose uptake at 5 mmol/gDCW/h, glucose uptake at 10 mmol/gDCW/h, glucose uptake at 10 mmol/gDCW/h with amino acid uptake constrained to 1 mmol/gDCW/h and glucose uptake at 10 mmol/gDCW/h with amino acid uptake constrained to 10 mmol/gDCW/h. In all conditions, oxygen uptake levels were kept at 5 mmol/gDCW/h. Each amino acid was evaluated individually under the corresponding supplementation conditions using flux sampling. Both OptGP (20) and ACHR sampling algorithms (21) were used to obtain flux distributions, generating a total of 10,000 flux samples using four parallel processes with a thinning factor of 1,000 for each sampling algorithm, enabling systematic comparison of their effects on MGSA flux. The resulting MGSA flux distributions were visualized using distribution plots. Among the amino acids screened, threonine and glycine emerged as strong candidates, showing a consistent effect on MGSA flux across the evaluated conditions (Figure 2 and Figure S1). Based on these findings, threonine was selected as our primary candidate for further investigation. To validate the robustness of this metabolic trend, we expanded our analysis across 56 *E. coli* genome scale metabolic models (GEMs) obtained from the BiGG database, the results of which are provided, with the mean MGSA flux summarized in (Figure 3).

**Figure 2.**
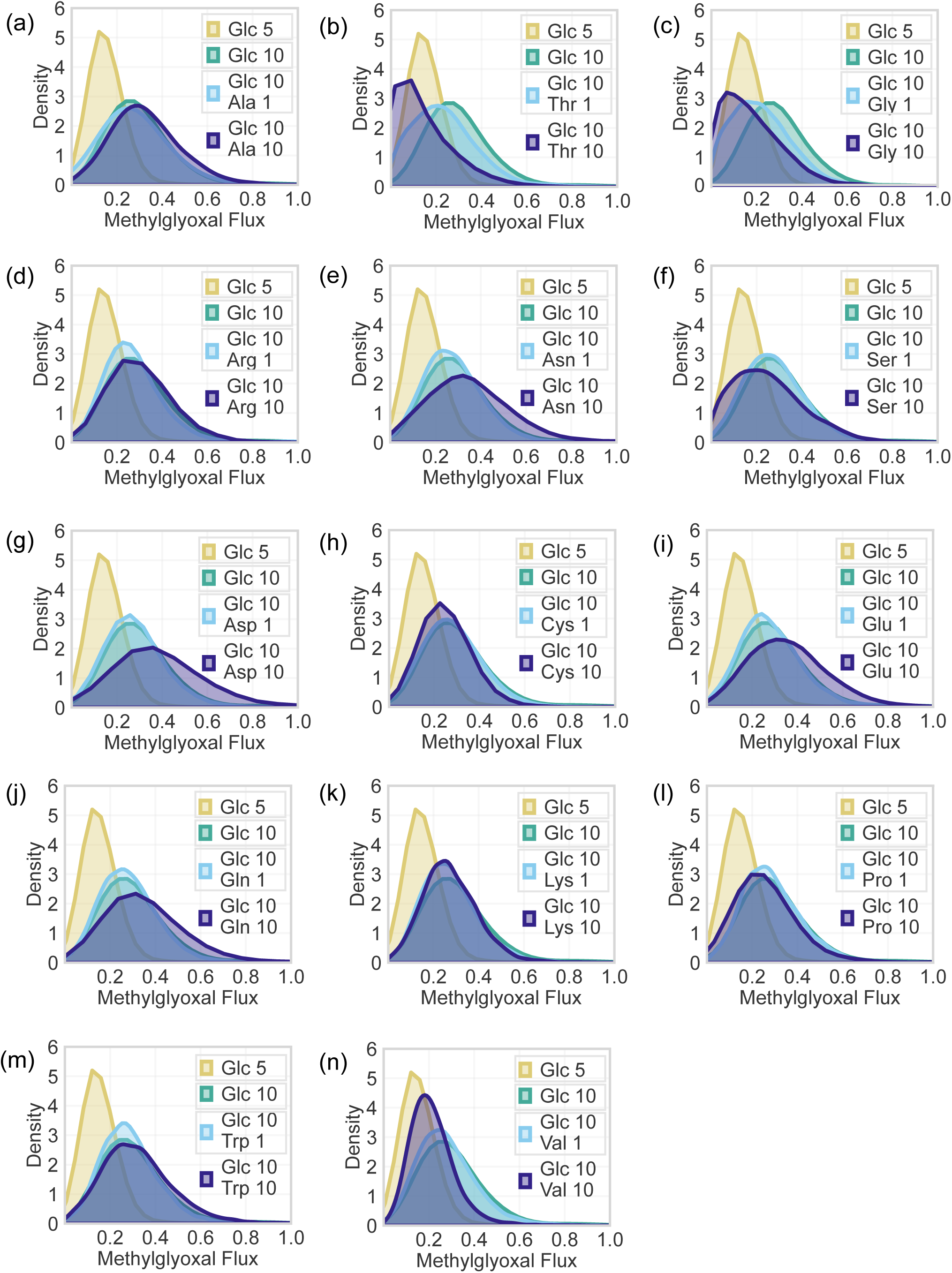
Identification of metabolites that modulate MGO flux distribution under varying glucose conditions and amino acid supplementation conditions performed using OptGp sampler. (Sampling in iJO1366 at 5 mmol/gDCW/h Oxygen uptake).

**Figure 3.**
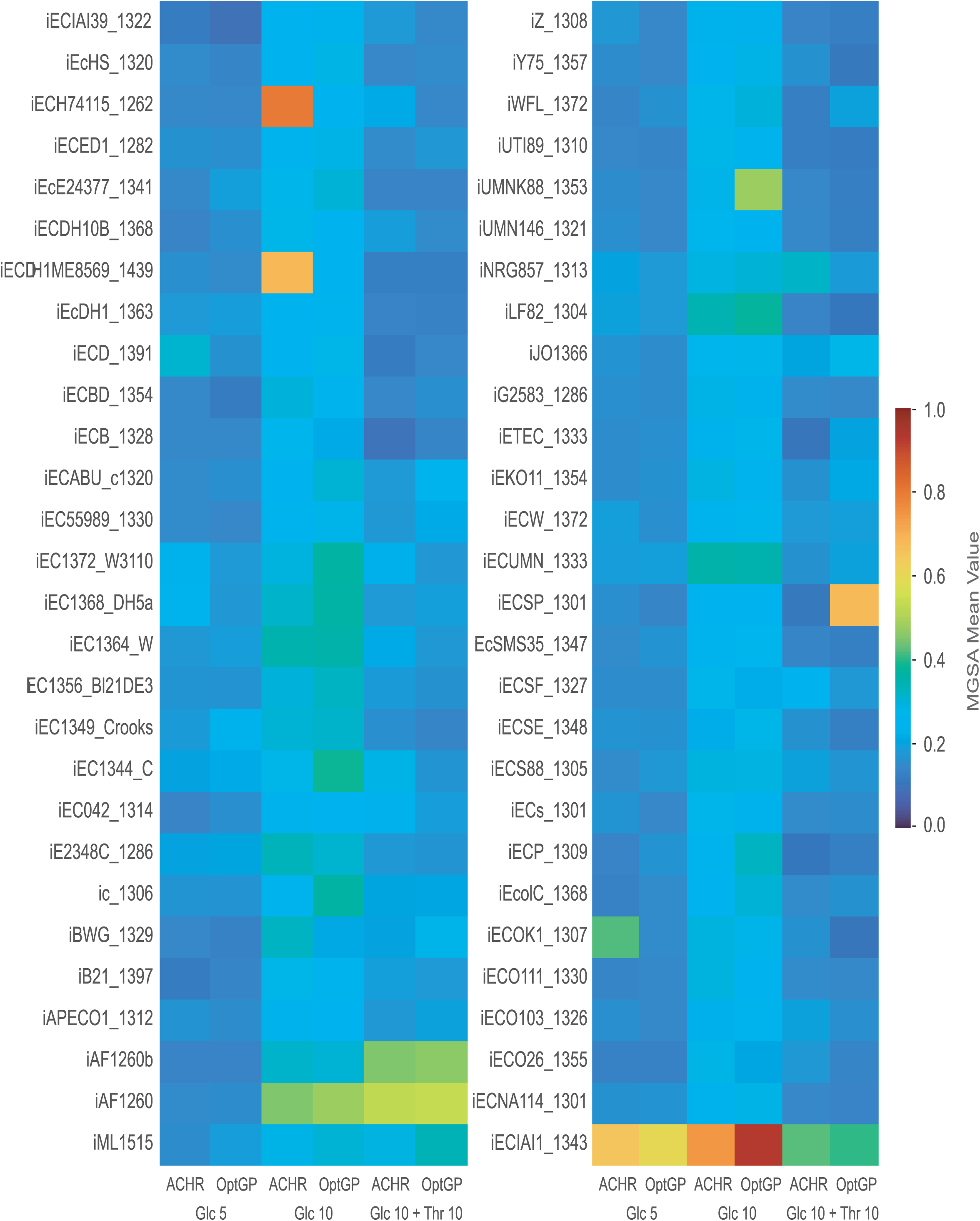
Mean MGSA flux (mmol/gDCW/h) across 56 *E. coli* genome scale metabolic models from the BiGG database. Each model was sampled using the ACHR and OptGP algorithms, generating 10,000 flux samples per condition. Simulations were performed under microaerobic constraints with the oxygen uptake lower bound fixed at 5 mmol/gDCW/h. Three nutrient environments were evaluated by setting the glucose uptake lower bound to 5 mmol/gDCW/h, 10 mmol/gDCW/h, and 10 mmol/gDCW/h supplemented with threonine at 10 mmol/gDCW/h.

#### 2.2.2 Flux sampling under varying oxygen uptake levels

All simulations were performed using the OptGP sampler, generating 10,000 flux samples per condition with a thinning interval of 250, 6 parallel chains, a burn-in of 2,000 iterations, and 10 computational processes. These parameters were selected to ensure robust exploration of the high dimensional feasible flux space. The large sample size stabilizes flux distribution estimates, thinning reduces autocorrelation, multiple chains improve mixing and mitigate chain specific bias, and the burn-in period removes transient initialization effects. Flux sampling followed the previously established protocol for threonine supplemented conditions, applied here across a graded oxygen uptake series spanning 1-10 mmol/gDCW/h in single unit increments. Flux samples from six OptGP chains were combined and annotated with chain and condition identifiers and the resulting ensembles were visualized using distribution based plots to capture feasible metabolic states and their condition dependent reorganization (Figure S2). To quantify how threonine supplementation alters the feasible flux space, we compared flux sampling distributions using two complementary metrics, the Kolmogorov-Smirnov (KS) distance and the Wasserstein distance. The KS distance captures the maximum divergence between cumulative distributions, providing sensitivity to structural changes in flux space such as the emergence or disappearance of feasible flux modes. The Wasserstein distance measures the average displacement required to transform one distribution into another, quantifying shifts in flux magnitude. High KS values indicate that threonine reshapes the architecture of the flux landscape, whereas high Wasserstein values reflect substantial changes in reaction flux intensities. Together, these metrics provide a rigorous statistical framework for assessing how threonine reorganizes metabolic behavior across oxygen environments.

### 2.3 Flux Balance Analysis

FBA was performed under the same conditions previously described in Section 2.2.2. The *in silico* analysis was performed using COBRApy (16) with the GLPK solver. FBA was performed with biomass as the objective to obtain the maximum predicted growth rate under the specified constraints. The biomass reaction was then constrained to maintain near optimal growth and MGO production was included as a secondary objective along with biomass.

## 3. Results

Using flux sampling and FBA we aimed to elucidate the metabolic reactions that can rewire MGO flux in bacteria. The flux sampling results were verified using 56 other *E. coli* genome scale models that revealed a highly consistent metabolic response under moderate microaerobic conditions, defined by an oxygen uptake rate of 5 mmol/gDCW/h (Figure 3). Across all models, MGSA flux increased when glucose uptake was raised from 5 to 10 mmol/gDCW/h, reflecting the expected metabolic shift toward enhanced MGO formation under elevated glycolytic throughput.

### 3.1 Identification of metabolites that modulate MGO flux

To identify the flux distribution of MGO across all feasible metabolic states, flux sampling analysis was performed under a relatively low glucose uptake rate (5 mmol/gDCW/h) and high glucose uptake rate (10 mmol/gDCW/h) with oxygen uptake fixed at 5 mmol/gDCW/h. The observations from these results revealed that MGO can exist across a range of 0.0001 (minimum flux at low glucose) to 4.553 mmol/gDCW/h (maximum flux at high glucose) (Table S1). Furthermore, flux sampling also identified the reactions that correlated with MGO flux. Top 10 reactions that are high when MGO flux is high, and top 10 reactions that are low when MGO flux is high are shown (Figure S3). The PCA plot that compared between high and low MGO flux states revealed that the overall metabolic state did not show a visible change even with high MGO in the *in silico* prediction model (Figure S4). Amino acids were supplemented into the high glucose condition (mean MGO flux 0.280) simulated model to evaluate its effect on MGO flux (Figure2, Figure S1). We observed threonine supplementation reduced mean MGO flux from 0.280 to 0.151 mmol/gDCW/h.

### 3.2 Threonine rescued glucose induced increase in MGO flux

It was seen that, threonine supplementation (10 mmol/gDCW/h) reduced MGSA flux in 53 of the 56 models, demonstrating a robust and broadly conserved inhibitory effect of threonine on MGSA production (Figure 3). Three models did not show MGSA reduction, a deviation attributable to model-specific differences in reaction content relevant to MGO metabolism (Figure 3).

For elucidating the mechanism behind threonine induced reduction of MGO in high glucose conditions, we performed FBA in the iJO1366 model. During low glucose (5 mmol/gDCW/h), the iJO1366 model gave a predicted flux value of 0.16 for MGO production (MGSA). When the glucose was increased (10 mmol/gDCW/h), MGSA flux showed a predicted increase to 1.87. By simulating the supplementation of threonine (10 mmol/gDCW/h) into the iJO1366 model, MGSA flux decreased from 1.87 to 0.405, even in the presence of high glucose. These predictions reconfirmed the role of glucose in increasing MGSA flux, in the simulation model. Furthermore, threonine was identified as a metabolite capable of rescuing the glucose induced increase in MGO.

### 3.3 Attenuating ALCD2x, GLYCL, GAPD and activating serine biosynthesis reactions reversed threonine dependent rescue of MGO flux

To identify the exact reasons behind the rescue effect of threonine, we tracked the bifurcating pathways downstream of threonine uptake and compared the flux values of these reactions using FBA. One of such reactions, ALCD2x which converts acetaldehyde to ethanol, showed an increase in flux from 0 (high glucose without threonine) to 7.73 (high glucose with threonine). When this reaction was inhibited by reducing the lower and upper bounds of ALCD2x to zero, threonine (10 mmol/gDCW/h) failed to partially reduce the MGSA flux, unlike before. When ALCD2x and GLYCL (bifurcating reactions downstream after threonine uptake) was blocked, along with minimal constraining of GAPD (from 18.2 to 17) and activation of serine biosynthesis PGCD (0 to 0.76), MGSA flux increased from 0.405 to 1.743. From these predictions we conclude that high glucose induced increase in MGO (1.87) was reduced by threonine supplementation at 10 mmol/gDCW/h. Threonine fails to reduce MGO when ALCD2x and GLYCL reactions are blocked, along with constraining GAPD and activating serine biosynthesis pathway. We also identified ALCD2x (acetaldehyde to ethanol conversion/NAD^+^ regeneration) as a major pathway through which threonine executes MGO reduction. The serine biosynthesis pathway suppression was also identified as a coordinated sub-mechanism.

### 3.4 Threonine supplementation increased flux at the payoff phase of glycolysis

To further understand the mechanism behind threonine induced MGO flux reduction, we studied the flux changes in the glycolysis pathway with and without threonine supplementation (10 mmol/gDCW/h) using FBA. In glycolysis, the reaction converting fructose 1,6 bisphosphate (F16BP) into dihydroxyacetone phosphate (DHAP) and glyceraldehyde 3-phosphate (GA3P) did not change much in the presence or absence of threonine supplementation (Figure 4). Unexpectedly we found that threonine supplementation controlled the flux from DHAP to GA3P (TPI) *in silico*. Threonine increased the flux of DHAP to GA3P from 7.67 to 9.01 followed by an increase in payoff phase fluxes, in this prediction model. downstream reactions from GA3P to pyruvate showed a corresponding increase in flux. These predictions underline the role of threonine in controlling the payoff phase of glycolysis in the *in silico* simulation model. Based on these *in silico* predicted observations, we hypothesize that increase in flux at the lower phase of glycolysis, might have reduced the stagnation of DHAP and overflow towards MGO. Therefore we hypothesize that threonine supplementation actively drives the ALCD2x pathway to provide localized NAD^+^ relief, which subsequently enhances the lower glycolytic flux (GAPD), thereby efficiently draining the accumulated DHAP pool and shutting down MGO production (Figure 5).

**Figure 4.**
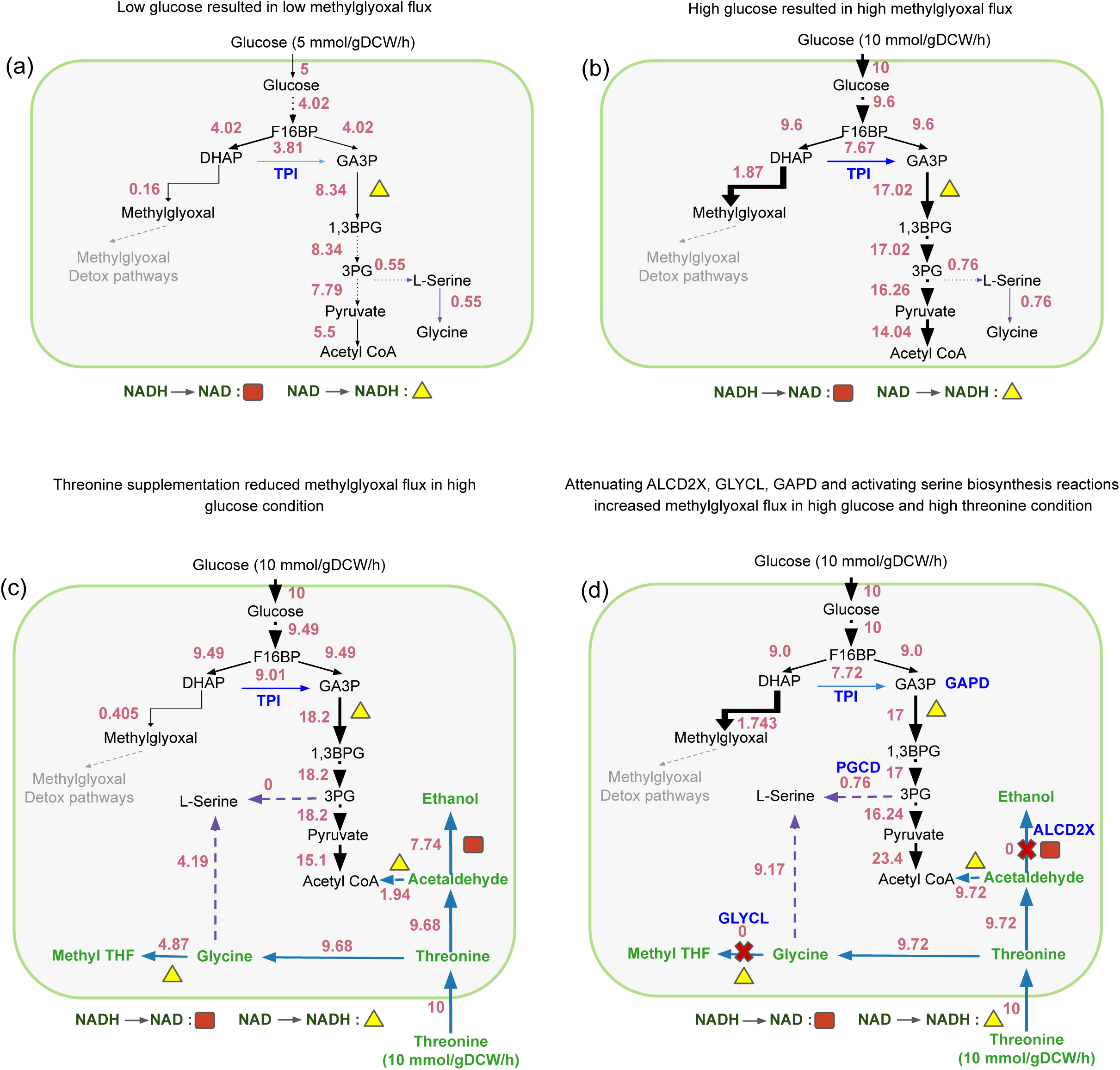
(a), (b), (c) Threonine rescued glucose induced increase in MGO flux. Overview of the mechanism involved in threonine mediated rescue of methylglyoxal flux in iJO1366 model at 5 mmol/gDCW/h microaerobic condition using FBA. (a) glucose uptake 5 mmol/gDCW/h, (b) glucose uptake 10 mmol/gDCW/h, (c) glucose uptake 10 mmol/gDCW/h and threonine 10 mmol/gDCW/h. (d) Attenuating ALCD2x, GLYCL, GAPD and activating serine biosynthesis reactions reversed threonine dependent rescue of MGO flux. Metabolic map depicting attenuated ALCD2x, GLYCL, GAPD and activated serine biosynthesis reactions in iJO1366 model at 5 mmol/gDCW/h microaerobic condition using FBA. Localized NAD^+^ metabolic rewiring in ALCD2x reaction acts as a major driver responsible for MGO reduction during threonine supplementation.

**Figure 5.**
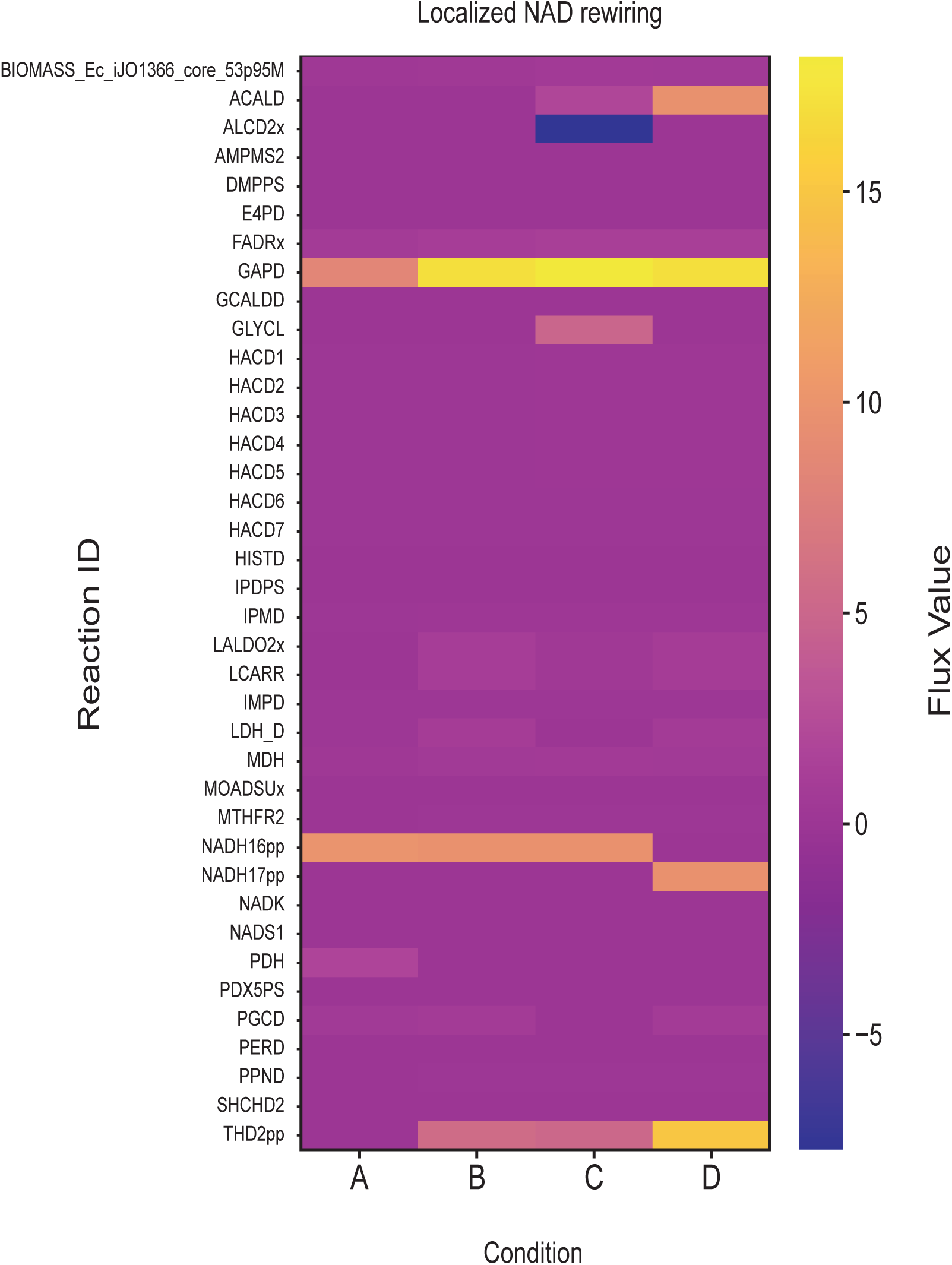
Heatmap representing all the active flux reactions involving NAD^+^/NADH, compared across all conditions using FBA. (Positive and negative flux values indicate flux in the forward and reverse directions, respectively). Condition A represents low glucose availability (5 mmol/gDCW/h), Condition B represents high glucose availability (10 mmol/gDCW/h), Condition C represents high glucose availability (10 mmol/gDCW/h) with threonine supplementation (10 mmol/gDCW/h), and Condition D represents the altered condition with attenuated ALCD2x, GLYCL and GAPD fluxes and activated serine biosynthesis pathway in high glucose (10 mmol/gDCW/h) and threonine supplemented condition (10 mmol/gDCW/h).

### 3.5 Glucose did not substantially increase MGO flux upon shifting to low oxygen conditions

We performed FBA under a low oxygen uptake value of 2 mmol/gDCW/h in the iJO1366 model at low (5 mmol/gDCW/h) and high (10 mmol/gDCW/h) glucose uptake rates. Unexpectedly, we observed that *in silico* glucose supplementation did not increase MGO flux to a noticeable unit (Figure 6). While MGO previously showed high flux (1.87) at high glucose (oxygen 5 mmol/gDCW/h), MGO flux reduced (0.247) upon shifting the model to low oxygen uptake levels (2 mmol/gDCW/h).

**Figure 6.**
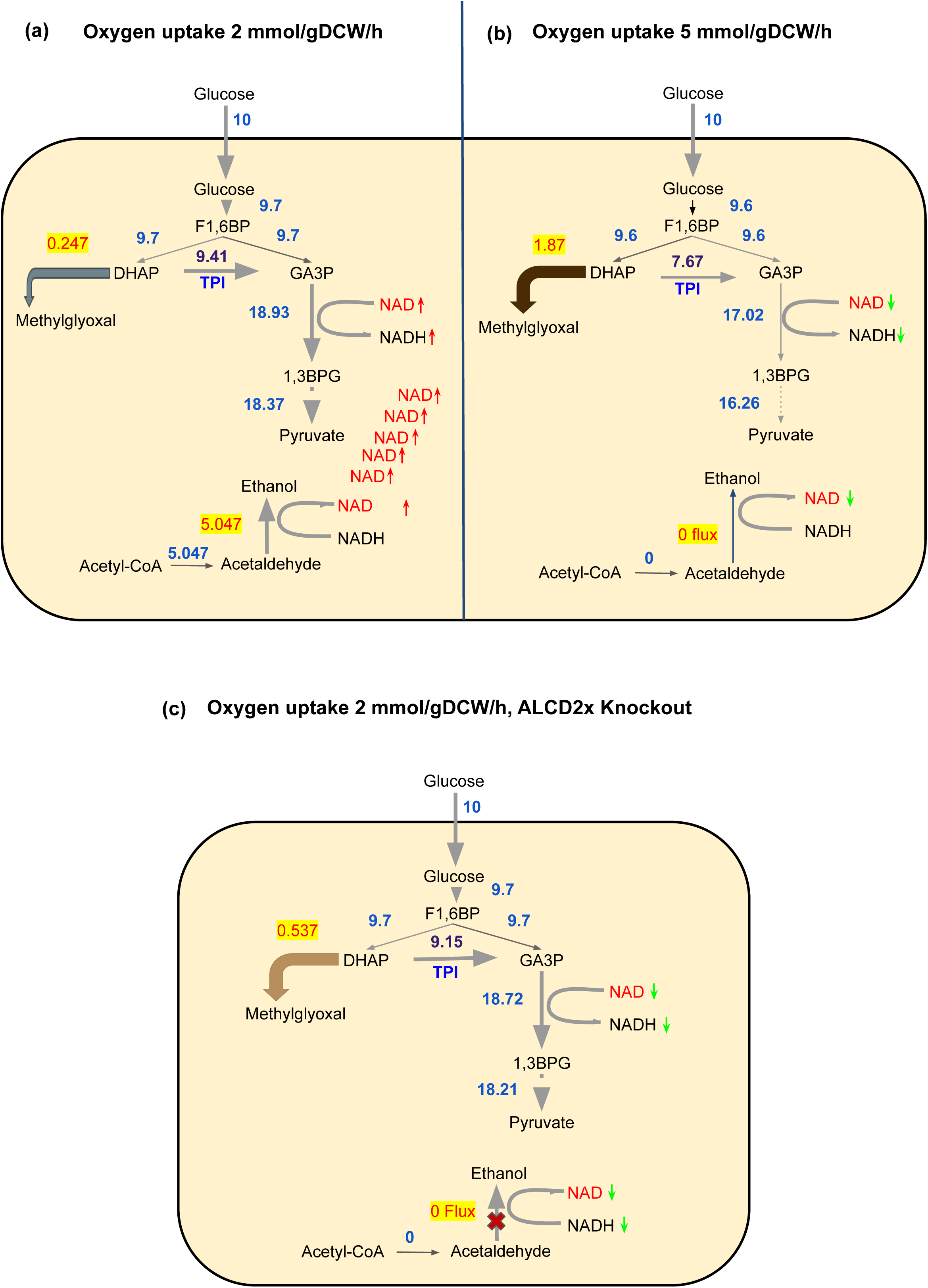
(a), (b) Glucose did not substantially increase MGO flux upon shifting to low oxygen conditions. Metabolic flux maps explaining the rewiring of NAD^+^/NADH mechanism via ALCD2x reaction in context to MGO flux production at 2 mmol/gDCW/h and 5 mmol/gDCW/h oxygen uptake levels using FBA. (c) Metabolic flux map explaining the partial increase in MGO flux upon knocking out ALCD2x at 2 mmol/gDCW/h oxygen uptake levels using FBA.

We tried to identify why MGO flux did not increase at 2 mmol/gDCW/h oxygen, even at high glucose (10 mmol/gDCW/h) in the model. Previous results showed that ALCD2x, which makes ethanol from acetaldehyde, has zero flux at high glucose conditions (oxygen 5 mmol/gDCW/h). Above results also confirmed that ALCD2x flux and ethanol production increased only after threonine addition. Unexpectedly, by shifting to low oxygen (2 mmol/gDCW/h), ALCD2x flux and ethanol production showed an increase, even without threonine supplementation, at high glucose conditions. We noticed that this increase in ALCD2x flux is due to an active pathway from acetyl-CoA to acetaldehyde which contributes to ethanol production during low oxygen uptake conditions (Figure 6). When we blocked ALCD2x at oxygen 2 mmol/gDCW/h, MGO levels partially increased from 0.247 to 0.537, which might be due to reduction in NAD^+^ regeneration, which also proves our hypothesis. (Figure 6). Based on these predicted observations, we arrive at the hypothesis that high glucose increased MGO at oxygen 5 mmol/gDCW/h, while high glucose failed to increase MGO at low oxygen 2 mmol/gDCW/h. We hypothesize that an increase in acetyl-CoA-acetaldehyde-ethanol flux during low oxygen, that regenerates NAD^+^, could be the mechanistic reason behind this observation, as blocking ALCD2x reversed these pathways and partially increased MGO flux.

### 3.6 Threonine failed to rescue glucose induced minimal increase in MGO during low oxygen conditions

While glucose addition brought only a minimal increase in MGO (0.147 to 0.247) at low oxygen (2 mmol/gDCW/h), we also observed that threonine could not reduce this minimal increase of MGO flux. Following this observation in order to study the dependency of MGO flux on oxygen, we performed a comparative analysis of varying oxygen uptake values of 1 mmol/gDCW/h to 10 mmol/gDCW/h under the conditions of varying glucose and threonine supplementation. The *in silico* simulation model predicted that MGO flux increases upon increasing oxygen uptake levels (Figure S5, Figure S6).

We also observed that even in oxygen uptake levels of 2 mmol/gDCW/h, threonine supplementation produces ethanol. Interestingly, ethanol production is more (11.7 mmol/gDCW/h) in 2 mmol/gDCW/h oxygen levels than in 5 mmol/gDCW/h oxygen levels (7.7 mmol/gDCW/h). Even after this high ALCD2x flux and NAD^+^ regeneration, threonine failed to reduce the very minimal increase of MGO caused due to glucose in lower oxygen levels (Figure 7). We hypothesize that this is because NAD^+^ generated from ethanol formation can control only DHAP to GA3P flux (glycolytic pull). NAD^+^ has no control over the minimal increase of MGO from 0.147 to 0.247 because this flux is contributed directly from glucose to DHAP to MGO route alone and the payoff phase of glycolysis. Therefore, threonine supplementation/NAD^+^ regeneration at 2 oxygen will fail to rescue 0.247 MGO flux, as this is not a stagnation dependent increase created as a consequence of insufficient NAD^+^ regeneration.

**Figure 7.**
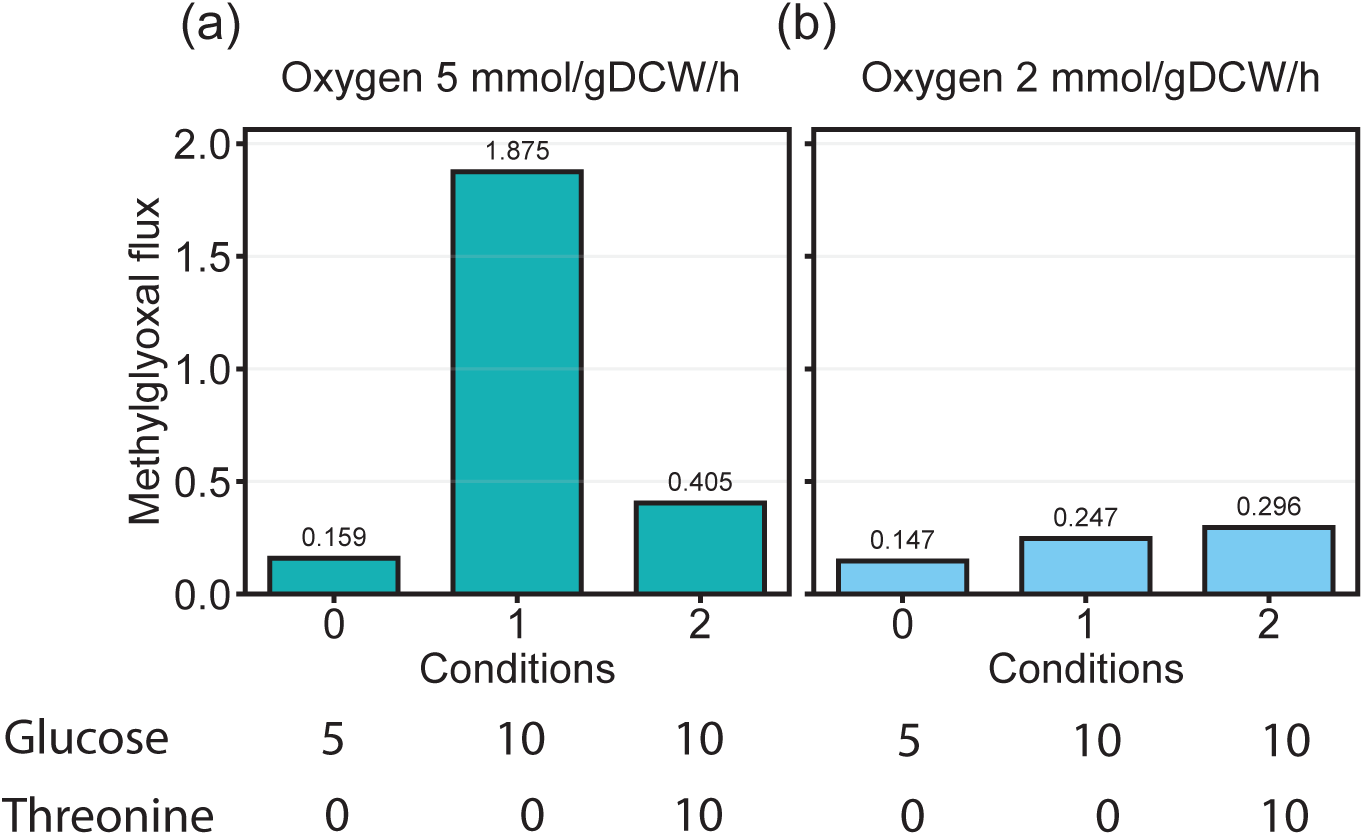
Bar plot representing oxygen dependent modulation of methylglyoxal flux in *E. coli* performed using FBA. MGSA flux was compared across glucose and threonine supplemented conditions under oxygen uptake constraints of 5 mmol/gDCW/h (left) and 2 mmol/gDCW/h (right). Condition 0 represents low glucose availability (5 mmol/gDCW/h), Condition 1 represents high glucose availability (10 mmol/gDCW/h) in the absence of threonine, and Condition 2 represents high glucose availability (10 mmol/gDCW/h) with threonine supplementation (10 mmol/gDCW/h).

## 4. Discussion

The excessive production of MGO during glycolytic overflow constitutes a major metabolic challenge for bacteria, yet how endogenous metabolic interventions prevent this overflow remains insufficiently understood. In this study, we performed FBA and flux sampling in the genome scale *E. coli* model iJO1366 to systematically screen for metabolic interventions capable of modulating glucose induced MGO overflow. Our major finding suggested that threonine supplementation significantly reduces glucose induced MGSA flux, by relieving the redox bottleneck at the payoff phase of glycolysis. This threonine induced NAD^+^ regeneration via ALCD2x maintains the glycolytic flux through GAPD, thus preventing DHAP accumulation and preventing subsequent conversion into MGO. This observation was consistent and well conserved across 53 of 56 independently curated *E. coli* genome scale models suggesting that it represents an intrinsic feature of *E. coli* central metabolism rather than a model specific artifact. Notably, only three models representing *E. coli* str. K-12 MG1655, consistent with iJO1366, deviates from this trend. Two models (iAF1260 and iAF1260b), which contain fewer reactions than iJO1366, show limited capacity to rewire flux under threonine supplementation. In contrast, the iML1515 model, which includes a larger reaction repertoire, shows only a modest increase. This behavior is likely driven by newly introduced ALR-group reactions that directly interact with the MGO metabolite. Our simulations also suggest that inactivation of the serine biosynthesis pathway during threonine supplementation, reflected by changes in PGCD and GLYCL, also influences MGO levels and contributes to its suppression. These findings suggest that MGO overflow represents a form of bacterial overflow metabolism, analogous to the acetate overflow mechanism in *E.coli* (22). Rather than being controlled primarily through direct regulation of methylglyoxal synthase, MGO production appears to emerge due to an imbalance between glycolytic carbon flux and the cellular capacity to maintain redox cofactor availability.

In *E. coli*, MGO formation is associated with accumulation of the triose phosphates dihydroxyacetone phosphate (DHAP) and glyceraldehyde 3-phosphate (GA3P), particularly when GAPDH activity is limiting or when carbon influx is excessive (23). *E. coli* is known to channel flux towards methylglyoxal upon accumulation of triose phosphates which results from phosphate depletion limiting the activity of GAP dehydrogenase or from excessive carbon intake (10). Our study further extends this established framework by introducing an upstream intervention in the form of threonine. This intervention relieves the imbalance without directly regulating the lower glycolysis or MGO detoxifying reactions. It is important to understand that threonine mediated rescue isn’t a common feature involved in MGO metabolic network, but a novel finding that is specific to the metabolic states under the constraints considered. Our result remains consistent in explaining the role of MGO as a glycolytic overflow metabolite whose production increases with enhanced glucose availability (24), rising from 5 to 10 mmol/gDCW/h, a pattern that aligns with clinical and metabolic observations in diabetes and other states characterized by an expanded intracellular triose pool driven by pathways such as gluconeogenesis and glyceroneogenesis (1, 13). Consistent with this relationship, glucose (or mannitol) as a preferred carbon source triggers Crh phosphorylation and reduces its binding to *MgsA*, indicating that MGO production is promoted when preferred carbon sources are available (25). We also understood how varying oxygen and glucose levels can affect the ability of threonine to suppress MGO production. At lower oxygen levels, glucose supplementation could not substantially increase MGO flux, also threonine supplementation did not further reduce it. This might be partially due to activation of ALCD2x under low oxygen levels which was inactive under moderate aerobic conditions.

During commercial scale production of a molecule, toxic byproducts accumulate. The energy expenditure of the cell to detoxify those toxic byproducts would eventually compromise the final yield (26). These byproducts beyond a tolerable threshold might in turn activate an energetically costly detoxification response and efflux pumping which would consume excess ATP and NADH (27). Such a circumstantial combination of large scale production along with increasing toxic metabolite stagnation and cell stress would result in a decline of production performance which is termed as a ‘’metabolic cliff’’ (28). This study demonstrates a general principle; Identifying a natural molecule that has a potential for redirecting an upstream precursor of a toxic overflow metabolite (e.g. MGO), before it crosses the tolerable thresholds, could bring a more energy efficient way to reduce toxic metabolite than depending on the downstream detoxification pathways.

Our discovery from the prediction models has its application in industrial production of MGO, as the natural molecules and reactions identified here can be utilized for fine tuning the commercial production and secretion of MGO. Manuka honey maintains its commercial success as an antibacterial agent due to its MGO abundance (29). However, the dependence on manuka honey for MGO brings enormous challenges; percentage of nectar collected specifically from manuka flower by the bees (30), prolonged time required for storage and maturation (non-enzymatic conversion of dihydroxyacetone to MGO) (31), environmental and seasonal variation in nectar yield (32) and a market value linked to MGO levels in the honey (30). Energetically favorable metabolic rewiring established in our study can be utilized to generate genetically engineered, MGO resistant bacteria capable of secreting MGO in large scale. Such engineered systems can simultaneously fine tune the MGO levels within the tolerable limits using the identified rescue metabolites, metabolic enzymes and oxygen conditions. This would enable a well controlled MGO synthesis from bacteria without any limitations like prolonged maturation time, environmental and seasonal variations materializing an economically viable, low cost supply chain.

While the results provide important insights, several limitations should be acknowledged. As FBA is a steady state modeling approach, it does not explicitly account for kinetics, allosteric regulation, or transcriptional control and therefore cannot capture the dynamic, real time states of the organism. Also these mechanisms might be well explained with respect to *E coli* strain K-12. Based on existing literature considering MGO as mainly the substrate produced from DHAP, other reactions producing MGO were not considered along with the secondary objective in FBA. This would have made the model highly constrained leading to infeasible states. Along with these limitations, PCA of the flux sampling data (Figure S4) showed substantial overlap between high and low MGO states, suggesting that differences in MGO flux do not substantially alter the overall metabolic flux distribution of the *in silico E. coli* model. Although experimental studies have demonstrated that elevated MGO causes macromolecular damage and can impair growth and cell survival (33, 34). These effects were not reflected in the global flux distribution of the *in silico E. coli* model, likely because the stoichiometric model does not explicitly represent MGO induced molecular damage. All the findings and predictions are model derived hypotheses and require direct experimentational validation.

## Supplementary figures

**Figure S1.**
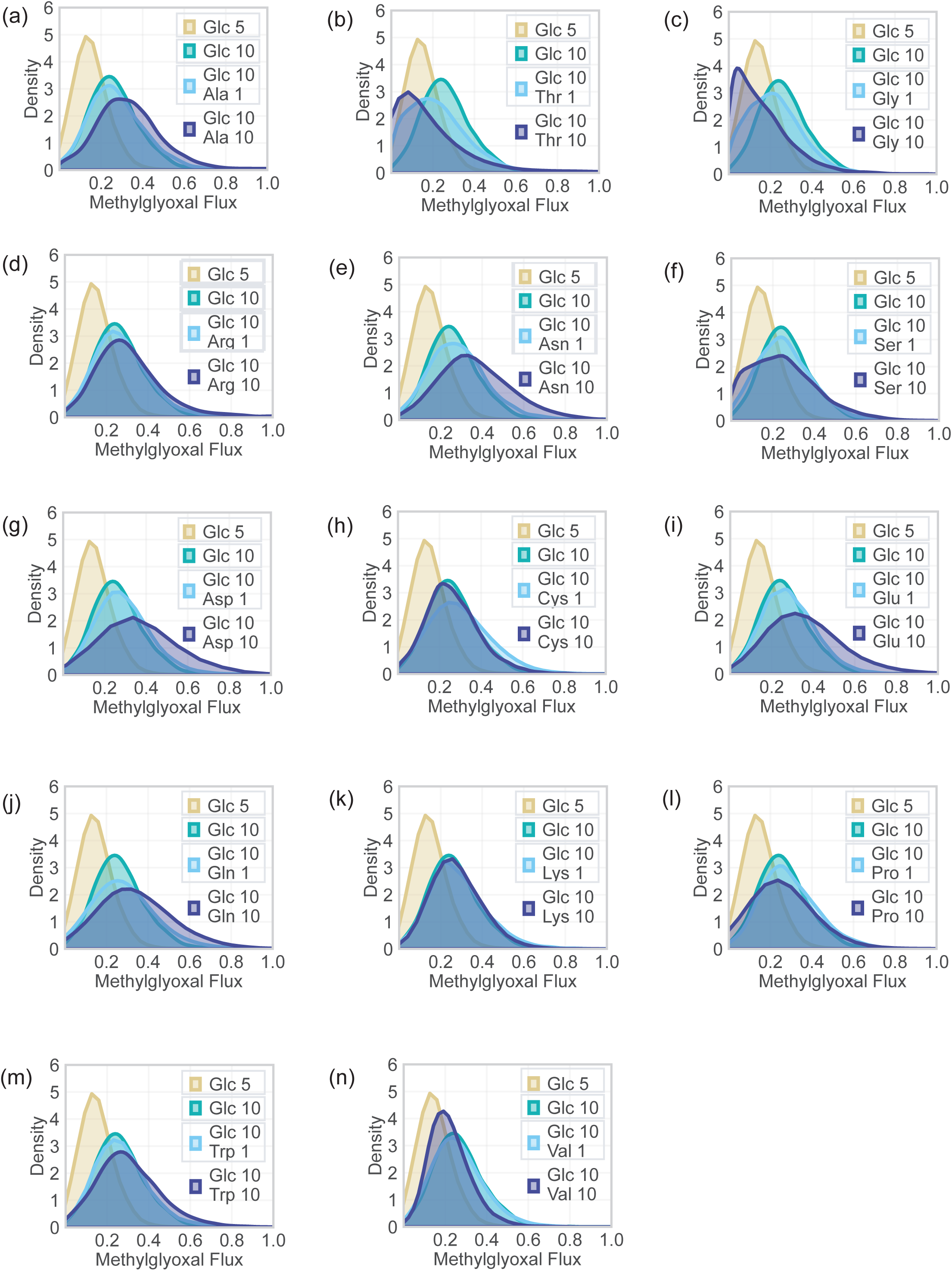
Identification of metabolites that modulate MGO flux distribution under varying glucose conditions and amino acid supplementation conditions performed using ACHR sampler. (Sampling in iJO1366 at 5 mmol/gDCW/h Oxygen uptake).

**Figure S2.**
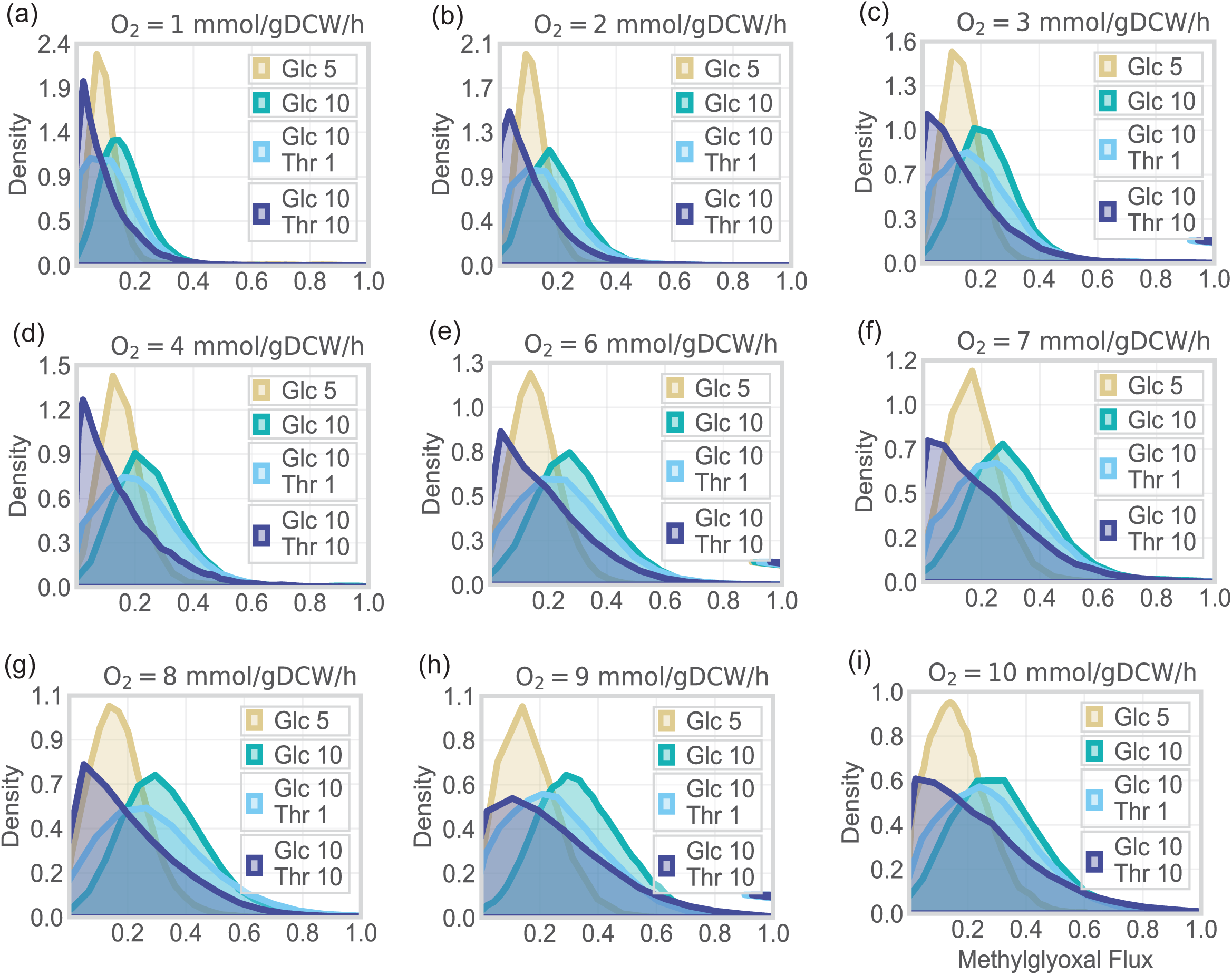
Distribution-based plots of MGSA flux (mmol/gDCW/h) generated by combining six OptGP sampling chains per condition. Four environmental conditions were analyzed across a graded oxygen uptake series spanning 1-10 mmol/gDCW/h in single-unit increments. The flux distribution corresponding to the 5 mmol/gDCW/h oxygen-uptake condition is presented in the main text (Figure 2).

**Figure S3.**
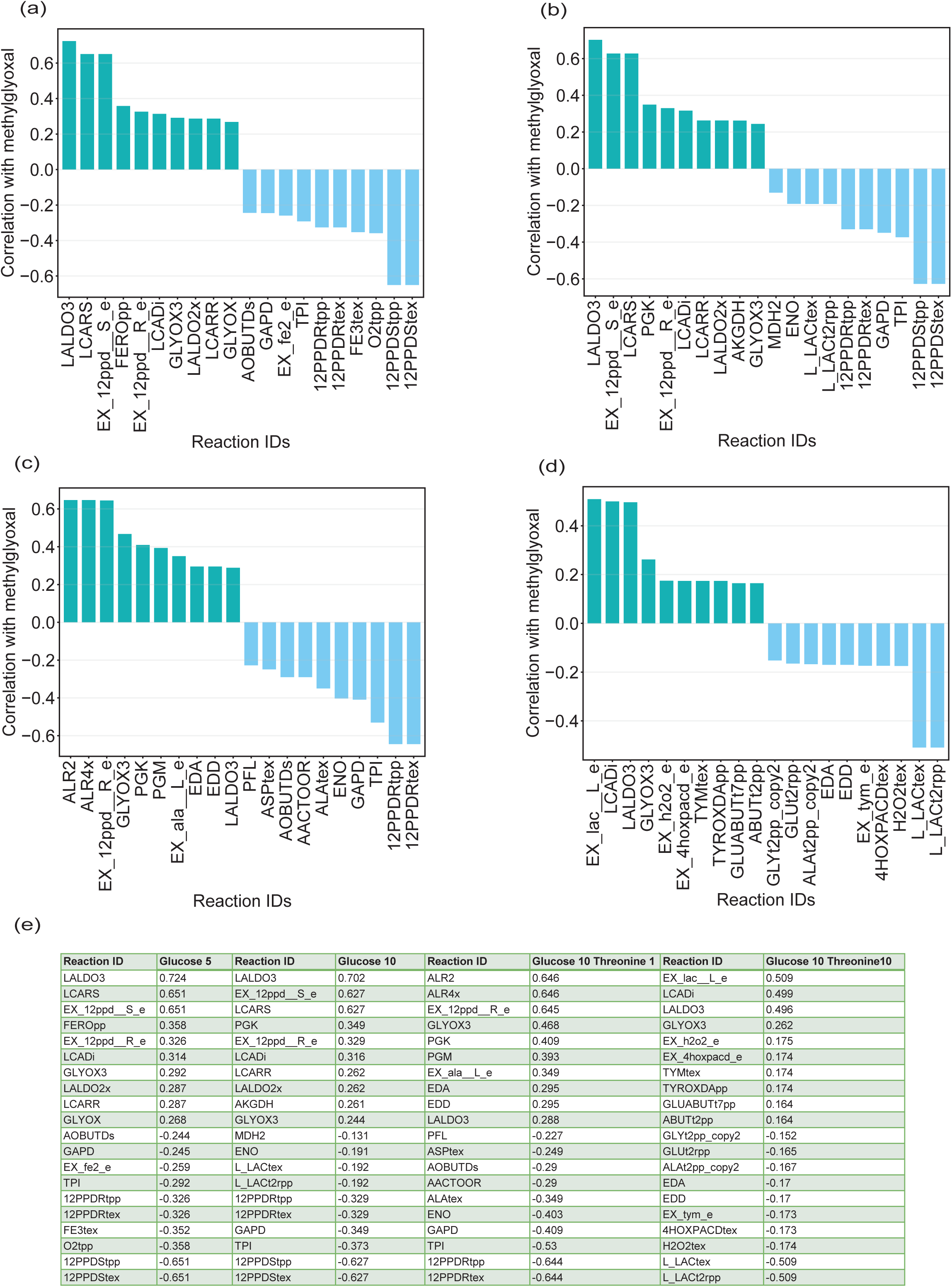
Correlation between flux through model reactions and methylglyoxal levels across different metabolic conditions obtained using flux sampling. (a), (b), (c), (d) represents bar plots showing pearson correlation coefficients between predicted reaction fluxes and methylglyoxal flux. (a) glucose uptake levels 5 mmol/gDCW/h, (b) glucose uptake levels 10 mmol/gDCW/h, (c) glucose uptake levels at 10 mmol/gDCW/h along with threonine uptake at 1 mmol/gDCW/h, (d) glucose uptake levels at 10 mmol/gDCW/h along with threonine uptake at 10 mmol/gDCW/h. Reaction IDs are shown on the x axis and correlation coefficients on the y axis. (e) Summary table listing the top positively and negatively correlated reactions for each condition, with corresponding reaction IDs and correlation coefficients.

**Figure S4.**
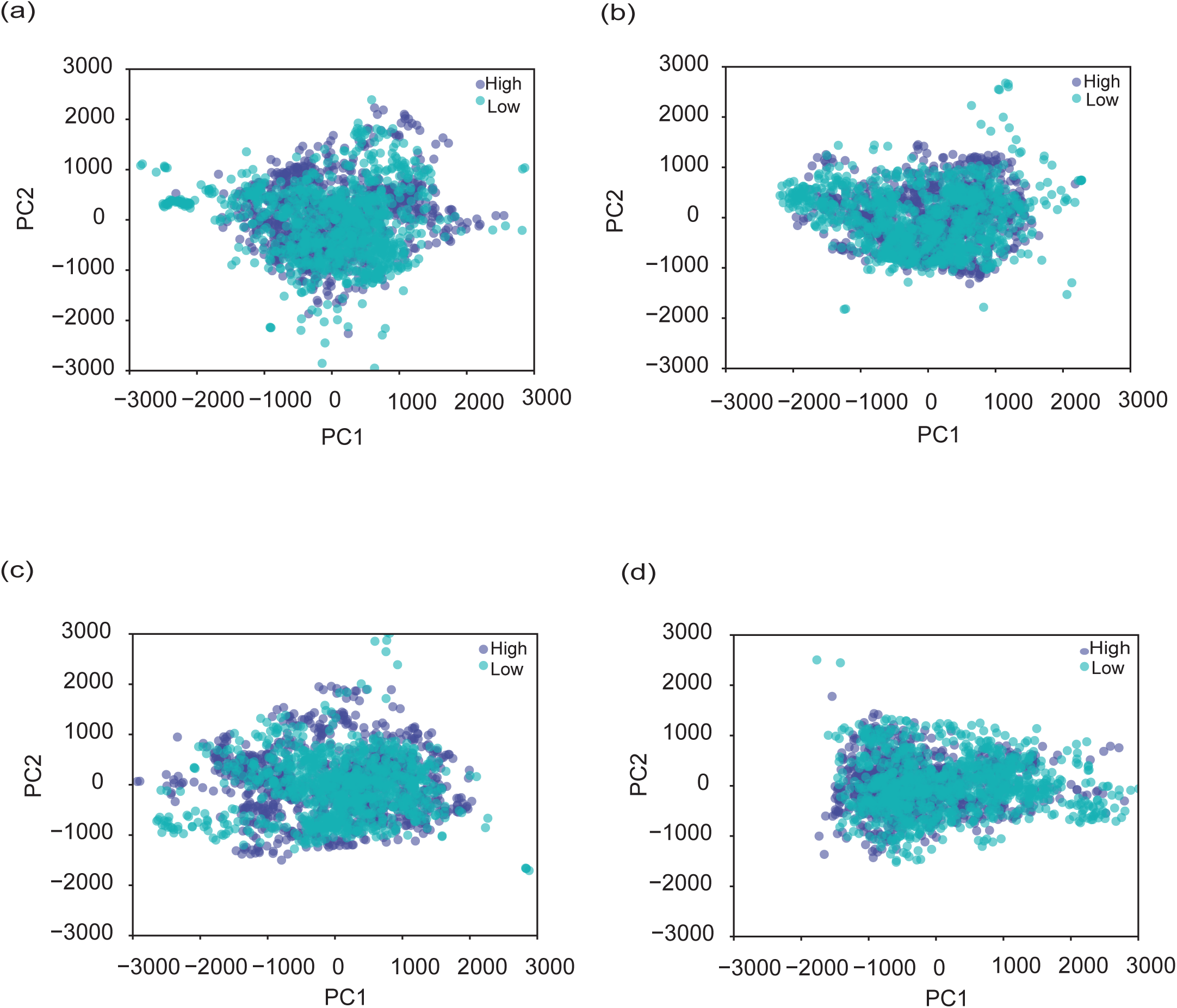
PCA of flux sampling distributions. Scatter plots of the first two principal components (PC1 and PC2) derived from flux sampling data under four culture conditions: (a) glucose 5 mmol/gDCW/h, (b) glucose 10 mmol/gDCW/h, (c) glucose 10 mmol/gDCW/h and threonine 1 mmol/gDCW/h, (d) glucose 10 mmol/gDCW/h and threonine 10 mmol/gDCW/h. Each point represents an individual sampled flux distribution, colored according to methylglyoxal levels, purple indicates high and teal indicates low methylglyoxal producing states.

**Figure S5.**
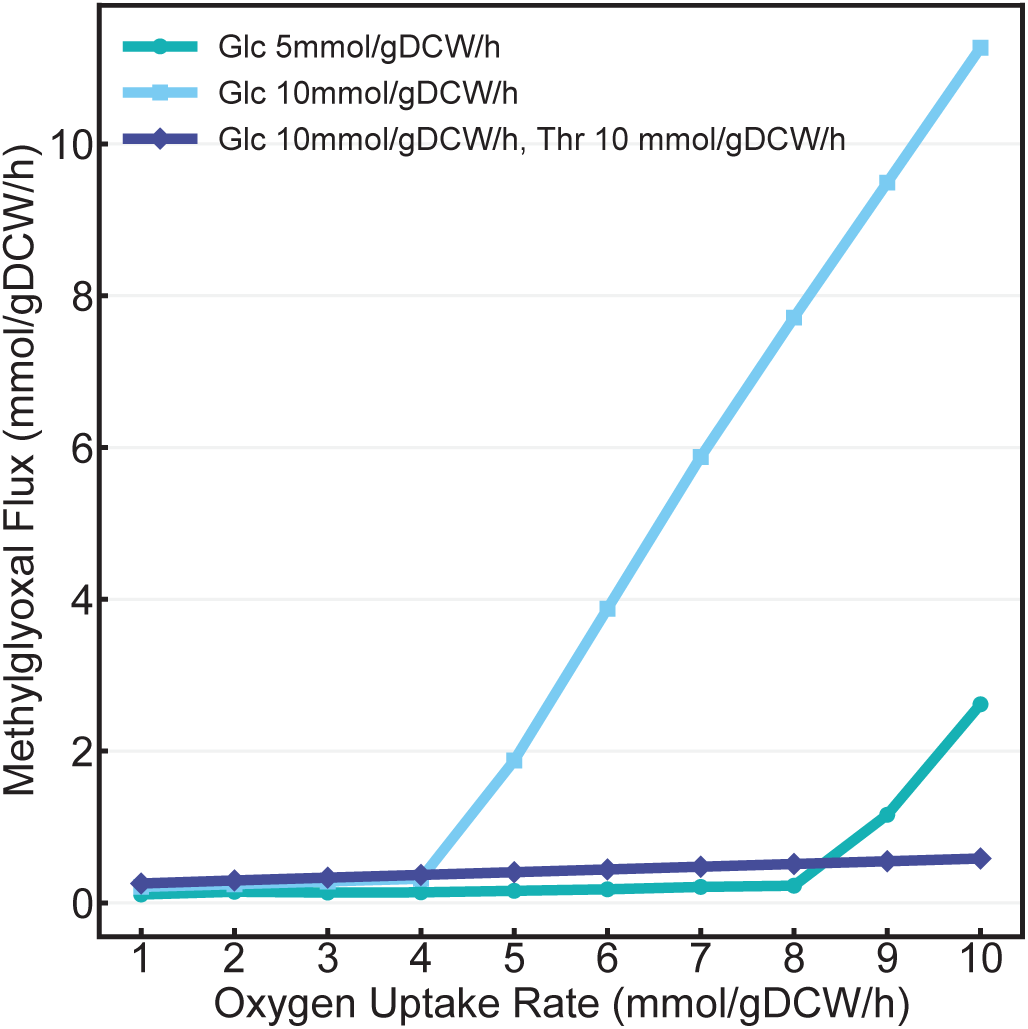
Line plot representing the effect of oxygen uptake rate on methylglyoxal flux under varying glucose and threonine supplemented conditions obtained using FBA.

**Figure S6.**
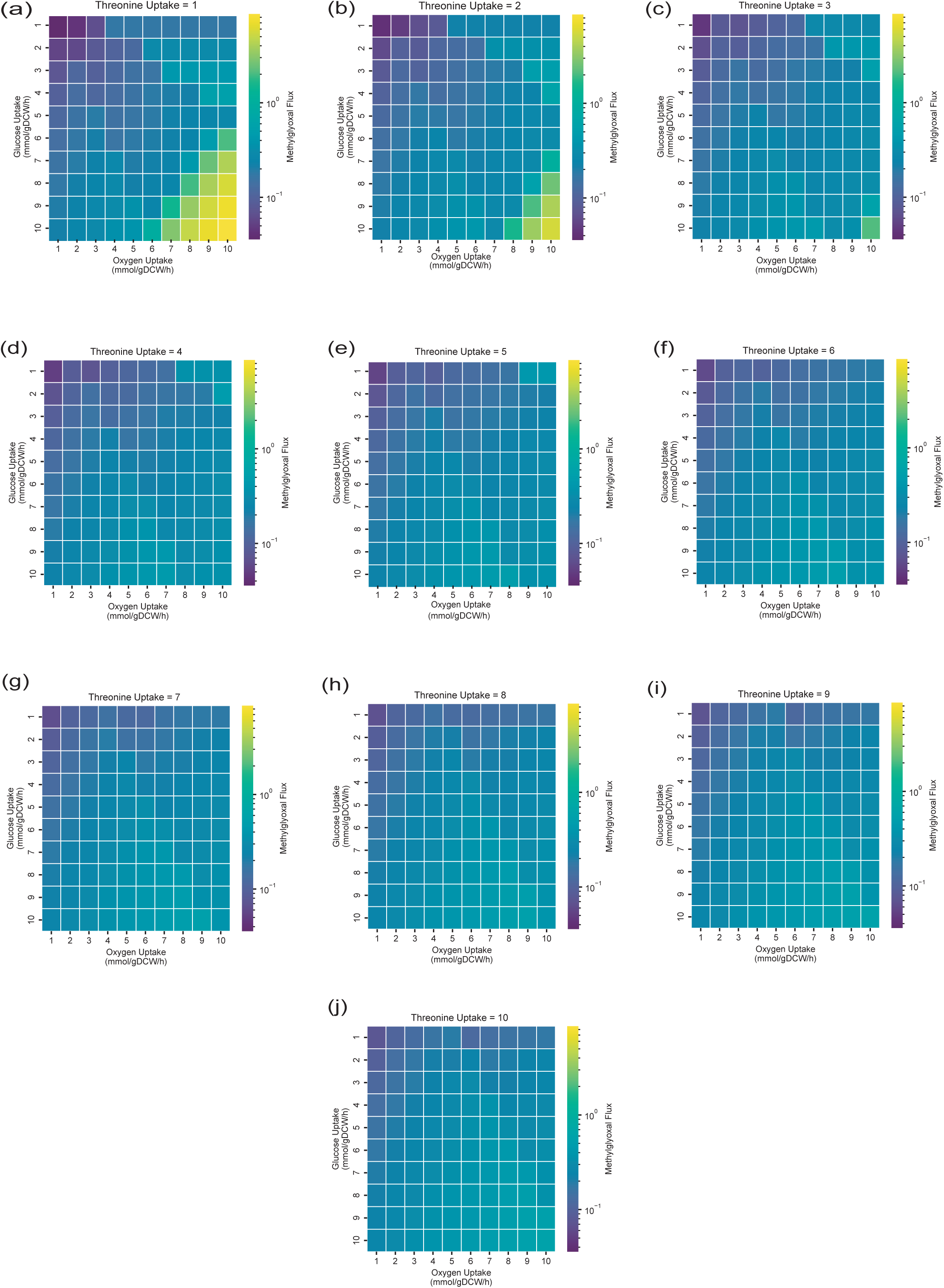
Three-way plot representing glucose, oxygen and threonine uptakes levels with respect to varying methylglyoxal flux across 1000 simulated FBA conditions. Heatmaps showing predicted methylglyoxal flux as a joint function of glucose uptake rate (y axis, mmol/gDCW/h) and oxygen uptake rate (x axis, mmol/gDCW/h), across a range of fixed threonine uptake rates from 1 to 10 mmol/gDCW/h (panels, as labeled, negative sign denotes uptake). Each panel represents a distinct threonine uptake constraint, with glucose and oxygen uptake systematically varied to generate 1,000 FBA solutions. Color intensity indicates the magnitude of methylglyoxal flux as indicated.

**Table S1.** Summary statistics of methylglyoxal levels across glucose and threonine supplemented conditions in Flux sampling.

| Conditions (mmol/gDCW/h) | samples | mean | std | min | 25% | 50% | 75% | max |
| --- | --- | --- | --- | --- | --- | --- | --- | --- |
| Glucose 5 | 10000 | 0.14845 | 0.099126 | 0.000134 | 0.093765 | 0.142686 | 0.194107 | 5.211848 |
| Glucose 10 | 10000 | 0.280326 | 0.154471 | 0.000547 | 0.188029 | 0.262744 | 0.354432 | 4.553046 |
| Glucose 10 and Threonine 1 | 10000 | 0.240625 | 0.182494 | 0.000062 | 0.134224 | 0.22111 | 0.315732 | 4.416346 |
| Glucose 10 and Threonine 10 | 10000 | 0.150522 | 0.172099 | 0.000014 | 0.045651 | 0.108043 | 0.211408 | 7.477098 |

